# Nucleosome positions alone determine micro-domains in yeast chromosomes

**DOI:** 10.1101/456202

**Authors:** O. Wiese, D. Marenduzzo, C. A. Brackley

**Affiliations:** SUPA, School of Physics and Astronomy, University of Edinburgh, Peter Guthrie Tait Road, Edinburgh EH9 3FD, United Kingdom

**Keywords:** chromatin domains, polymer simulations, MicroC

## Abstract

We use molecular dynamics simulations based on publicly available MNase-seq data for nucleosome positions to predict the 3-D structure of chromatin in the yeast genome. Our main aim is to shed light on the mechanism underlying the formation of micro-domains, chromosome regions of around 0.5-10 kbp which show enriched self-interactions, which were experimentally observed in recent MicroC experiments. We show that the sole input of nucleosome positioning data is already sufficient to determine the patterns of chromatin interactions and domain boundaries seen experimentally to a high degree of accuracy. Since the nucleosome spacing so strongly affects the larger-scale domain structure, we next examine the genome-wide linker-length distribution in more detail, finding that it is highly irregular, and varies in different genomic regions such as gene bodies, promoters, and active and inactive genes. Finally we use our simple simulation model to characterise in more detail how irregular nucleosome spacing may affect local chromatin structure.

## Introduction

Recent advances in next-generation sequencing (NGS) technologies have revolutionised our understanding of how the spatial organisation of genomes within the cell nucleus impacts on gene regulation and cell function. Specifically, the chromosome conformation capture (3C) family of methods give information about interactions between different chromosome regions (i.e., within a population of cells, how likely are two loci to be spatially proximate). High-throughput variants of the method such as Hi-C have shown that most genomes (ranging from bacteria [1] to mammals [2, 3]) are organised into domains, where regions within the same domain are more likely to interact with each other than with regions in different domains. These are usually known as either chromosomal interaction domains (CIDs) or topologically associated domains (TADs). As the sequencing depth of HiC data has increased, allowing interactions to be probed at higher resolutions, domains have been found at many different length scales. In mammals, the highest resolution data has revealed TADs ranging in size from 40 kbp–3 Mbp [4], and analysis of interactions between neighbouring TADs revealed cell specific “metaTADs” [5]; this points to a hierarchical domain structure [6], with domains observed at many scales. The functional role of domains is only just beginning to be understood: domain boundaries have been shown to provide insulation between enhancers and promoters, which is particularly important for developmental genes [7]; disruption of boundaries can lead to mis-regulation of genes [8]; and large scale re-arrangement of TADs has been implicated in diseases such as cancer [9].

Recently a new genome-wide chromosome conformation capture method called MicroC, developed by Hsieh *et al.* and detailed in Refs [10, 11], has allowed chromatin interactions to be probed at the nucleosome level. The technique uses a protocol where chromatin fragmentation is achieved by micrococcal nuclease (MNase) digestion, and has yielded nucleosome resolution interaction maps of the entire genome of the budding yeast *Saccharomyces cerevisiae*. At this higher resolution, Hsieh *et al.* were able to identify domains with sizes between 0.5–10 kbp or 4–50 nucleosomes. These typically contain between zero and eight genes, and their boundaries are associated with nucleosome depleted regions (NDRs; often found at gene promoters) as well as enrichment of histone modifications associated with transcriptional activity, chromatin remodelling factors and the cohesin loading factor Scc2 [10]. Recent conventional HiC experiments on budding yeast [12] also revealed a domain structure at a larger scale, with an average size of 200 kbp. The boundaries of these larger domains are enriched in transcriptional activity, and seem to be strongly linked to replication timing. The smaller sized domains thus appear to be a distinct level of organisation – here for clarity we will call these “micro-domains”.

In this paper we use computer simulations based on polymer models to study the formation of nucleosome level micro-domains in yeast. Our aim is to understand how these domains are formed, and what determines their boundaries. In other words, we want to understand the essential model ingredients which are required to yield the domain patterns observed in the MicroC data of Ref. [10]^1^. Intriguingly, we discover that a deceivingly simple polymer model which includes only the average nucleosome positions as an input can already predict many features of the 3-D organisation – most notably the locations of micro-domain boundaries – to a high degree of accuracy. Surprisingly, we find that a model with more realistic DNA-nucleosome geometry does not in fact show significant differences, or improved agreement with the data. This suggests that the information which encodes for 3-D micro-domain structure is already present within the map of nucleosome positions. More specifically, we find that it is the irregular spacing of nucleosomes in yeast chromatin which leads to boundary formation. Although nucleosome mapping data showing this irregular spacing have been available for some time, the textbook picture of a regular fibre is still prevalent. To better understand how nucleosome spacing genome-wide differs from a regular fibre we examine the distribution of linker lengths at micro-domain boundaries, and in different genomic environments (i.e. with active and inactive genes). We find that the linker length distribution shows peaks at short, medium and long ranges, and these are distributed differently in active and inactive regions. By analysing our simulated chromatin structures, we find that the local compaction of fibres with irregular spacing, such as those constituting the yeast genome, is highly heterogeneous, and very much unlike that of regular fibres such as those reconstituted *in vitro*.

## Results

### A simple nucleosome-level model for chromatin.

We start with the simplest possible model for chromatin which resolves individual nucleosomes and linker DNA. Using a beadand-spring polymer modelling approach, DNA is represented as a semi-flexible polymer where 2.5 nm beads correspond to approximately 8 bp of DNA. This is a well studied model [16–18], and uses simple phenomenological interaction potentials to give the correct bending rigidity for DNA *in vivo*. Specifically the DNA has a persistence length *l_P_* = 50 nm (this is a measure of the bending stiffness, defined as the distance along the molecule over which correlations in backbone orientation decay). Nucleosomes (including both histone proteins and wrapped DNA) are represented by 10 nm beads. A chromatin fibre is then modelled as sections of DNA (linkers) interspersed with nucleosome beads. For simplicity we do not include any orientational or bending constraints between linkers and nucleosomes: i.e., where there is a connected chain of DNA-nucleosome-DNA beads this acts as a freely rotating joint. We do not include any interactions between nucleosomes, or between nucleosome and DNA, other than simple excluded volume. A schematic of the model is shown in Fig. 1a, and full details of all interaction potentials and parameters are given in the Supplementary Material. Note that a similar model was recently proposed [19] which represented nucleosomes as spheres, but did not include linker DNA. More detailed nucleosome-level models of chromatin have been studied in the literature [20–22], but these are much more computationally expensive, and so are limited to short fibres; our approach here – starting with the simplest possible model and adding details incrementally – allows us to understand which aspects of the model are important for the resulting behaviour.

**Fig. 1:**
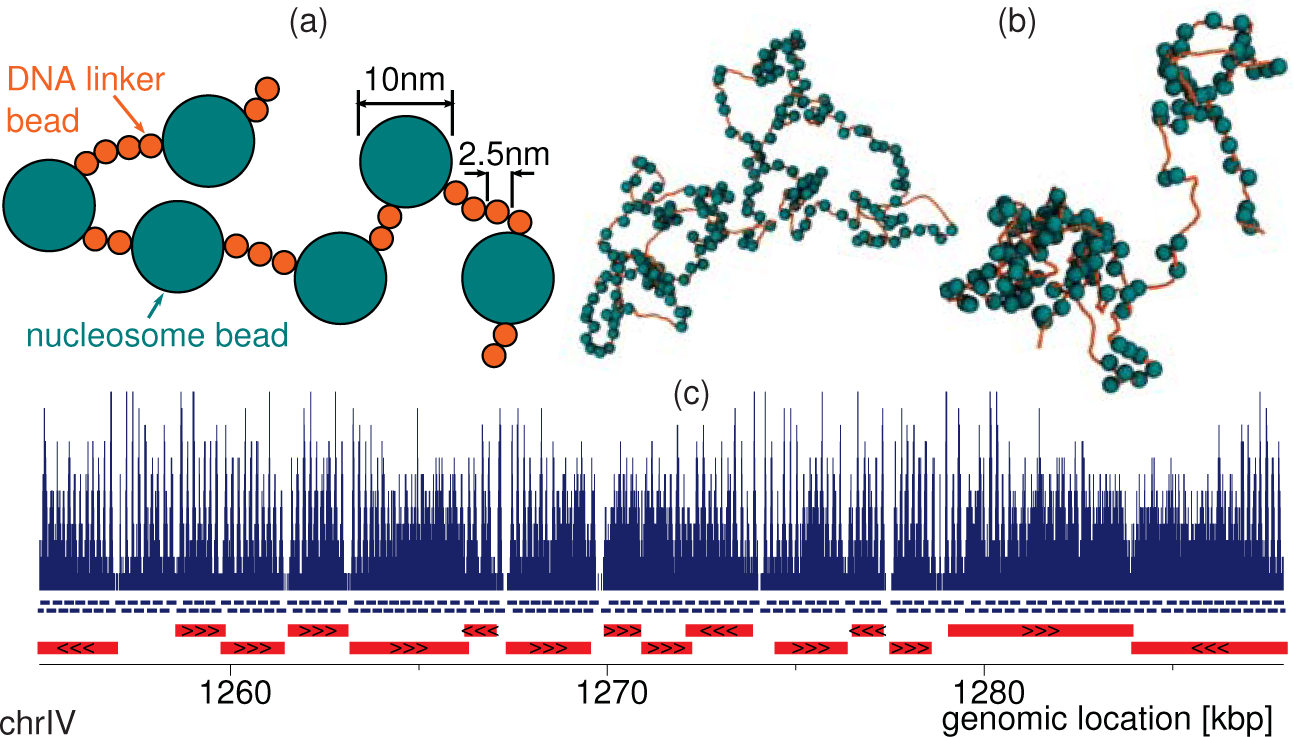
A simple chromatin model using nucleosome positions as input. (**a)** Schematic showing the bead-spring-polymer model for a chromatin fibre. DNA is represented as a chain of 2.5 nm beads connected by springs, and including a bending rigidity to give a persistence length *1_p_* = 50 nm. Nucleosomes (including both histone proteins and wrapped DNA) are represented by 10 nm spheres. (**b)** Typical snapshots from simulations (shown at different zoom levels). Left: simulation of region chrIV:1254937-1287938 of the yeast genome (SacCer3 build). Right: simulation of region chrXI:86225-108599. (**c)** Data used as an input to the model. Top: pile-up of reads from yeast MNase data (from Ref. [13]) for the indicated genomic region. Blue lines under the plot show the positions of nucleosomes determined using the NucPosSimulator software [14]. Red bars show genes. Similar plots for other genomic regions are shown in Suppl. Fig. S1.

In order to simulate a specific genomic region, we use MNaseseq data [13] to infer the most likely nucleosome positions and linker lengths. Importantly this is the only data which is used as an input to the model. Like the MicroC protocol, these experiments use micrococcal nuclease to digest any DNA which is not protected by nucleosomes, but rather than interaction data they provide a genome-wide map showing nucleosome coverage within a population of cells (Fig. 1c top). In order to infer the most likely positions of nucleosomes from this data we use the NucPosSimulator software [14]. In short, this uses a simulated annealing protocol to position nucleosomes according to an effective potential which is obtained from the MNase-seq data – further details are given in the Supplementary Material (see also Ref. [14]). In Fig. 1c nucleosome positions are shown with blue lines under the plot.

We use the LAMMPS molecular dynamics software [23] to perform Langvin dynamics simulations (see the Supplementary Material for full details). We selected 8 regions of between 15-43 kbp long across six different yeast chromosomes in order to get a representative sample of genic chromatin regions. In total our simulations covered ~ 240 kbp. After suitable equilibration, we evolved the dynamics to obtain a set of chromatin conformations for each region. (Typically after a 122 *τ* equilibration simulation, we simulated for a further 50 × 10^3^ *τ* and saved configurations every 250 *τ*. This was repeated 20 times for each region, resulting in a population of 2000 configurations per region. Here *τ* is the simulation time unit, equivalent to 80 *µ*s – see the Supplementary Material for full details.) Then, from each population of simulated chromatin conformations we generate a simulated MicroC map. Specifically this generates a map of *nucleosome-nucleosome* interactions; in order to compare with MicroC data from Ref. [10], we took that data and mapped each interaction to a specific pair of nucleosomes using the same nucleosome positions as in the simulations. Note that this means that in figures interactions are shown at a nucleosome level, and not in base-pair coordinates (as is common in HiC).

### Nucleosome spacing alone is sufficient to determine chromatin micro-domains.

Though this model is simple, as it treats nucleosomes as spheres, rather than a more realistic disk-like shape, and it ignores the complex inter-nucleosome interactions mediated by histone tails, surprisingly we find that it captures sufficient detail to correctly predict many features of short-range nucleosome contacts in 3-D.

Figure 2a shows results from a simulation of a 33 kbp region of the yeast genome (chrIV:1254937-1287938); a snapshot of a typical conformation for this regions is shown to the left in Fig. 1b. The nucleosome-nucleosome interaction map shows simulation results in the upper left half and the corresponding MicroC data from Ref. [10] in the lower right (the simulated map is constructed such that the total number of reads is the same as in the data). First, we note the striking visual similarity between the two maps, especially close to the diagonal. Second, to more quantitatively compare the simulations with the experiments, we identified micro-domains by calling boundaries (see the Supplementary Material for details). In this region the MicroC data shows 17 boundaries; remarkably our simulations correctly predict the location of 13 of these (76%). We identify boundaries by fist calculating a “boundary signal” for each nucleosome (see the Supplementary Material); comparing the simulated and MicroC boundary signals we find a correlation coefficient *r* = 0.50 (*p* < 10^−10^ using the Spearman Rank correlation). As well as the correctly predicted boundaries, the simulations also predict an additional 12 boundaries which are not found experimentally. Extending this analysis to all 8 simulated regions, which cover a total of 240 kbp (Fig. 2b), we find that the simulations correctly predict the positions of 84.5% boundaries (93 out of 110), but also predicted 58 additional boundaries (i.e 61.5% of simulation boundaries were correct), and the correlation coefficient for the boundary signal is *r* = 0.52 (*p* < 10^−10^). Since the only input data to the model is nucleosome positions, we conclude that this is a major driver of chromatin interactions at this scale. One might expect that a pair of widely spaced nucleosomes could act as a boundary to nucleosome interactions. Indeed the nucleosome spacing (or linker lengths) at boundaries tends to be much larger than average (within the simulated regions boundary linkers are on average ~ 120 bp, compared to ~ 28 bp for all linkers; see Suppl. Fig. S3a).

**Fig. 2:**
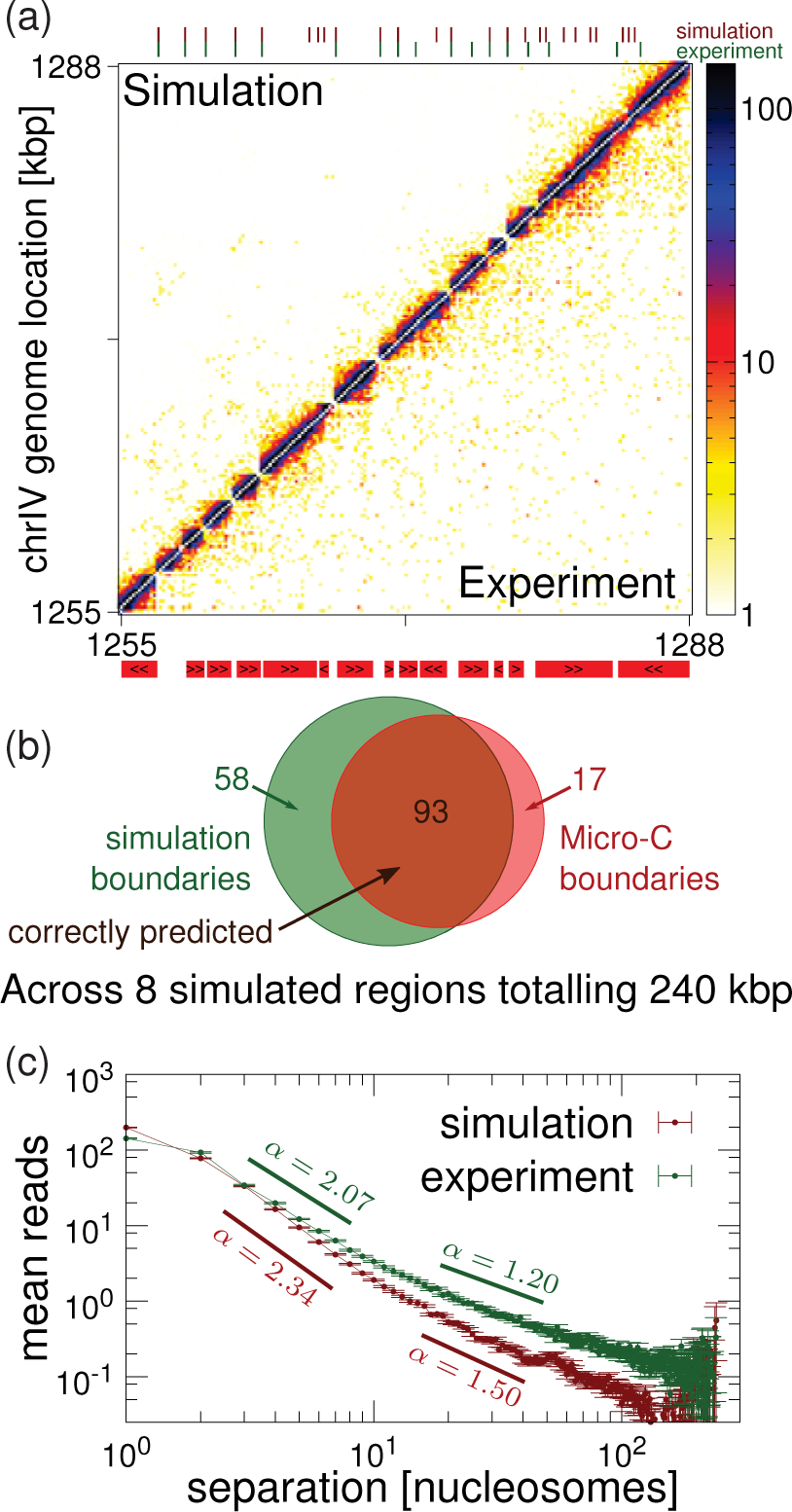
Chromatin fibre simulations accurately predict MicroC interactions for *Saccharomyces cerevisiae*. (**a)** Map showing interactions between nucleosomes in region chrIV:1254937-1287938. The colour of the square at coordinates *i*, *j* indicates the number of MicroC reads corresponding to interactions between nucleosomes *i* and *j*. The lower triangle shows MicroC data from Ref. [10], while the upper triangle shows simulation results. Numbers on the axes show genomic position, but this is approximate, since nucleosome spacing is not regular. The locations of genes (mapped to the nearest nucleosomes) are shown below the plot (red bars; gene orientation is indicated with black arrowheads). Domain boundaries were called from each map (see the Supplementary Material for details), and these are indicated with ticks above the plot (upper row shows simulation boundaries in red, lower row shows experimental boundaries in green). Similar plots for all 8 simulated regions are shown in Suppl. Fig. S2, with the MNase data used as input in Suppl. Fig. S1. (**b)** Venn diagram showing the number of domain boundaries across all 8 simulated regions. Overall ~ 84.5% of boundaries were correctly predicted by the simulations; an additional 58 boundaries not present in the data were found in the simulated interaction maps. A “correct prediction” is defined as a boundary in the simulated map being within one nucleosome of a boundary in the experimental map. (**c)** Plot showing how, on average, the number of interaction reads between nucleosomes scales with their genomic separation (measured in nucleosomes, i.e. a separation of 1 means neighbouring nucleosomes).

Examining the boundaries found in the simulations in more detail, we find that the 17 “missing” boundaries (i.e. those present in the MicroC data but not found in simulations) tend to be at more closely spaced nucleosomes (short linkers, on average ~ 22 bp; Suppl. Fig. S3b). Using data for histone modifications and protein binding (from Refs. [24] and [25] respectively) we find that the correctly predicted boundaries tend to be flanked by nucleosomes enriched in marks associate with gene activation (H3K9, H3K18 and H3K56 acetylation and H3K4me3; Suppl. Fig. S3c) consistent with the findings of Ref. [10]. Interestingly, the “missing” boundaries lack any significant enrichment of these marks (i.e. they do not display the features found at most boundaries), and they tend to be weaker (see the Supplementary Material for details of boundary strength quantification). Together this points to the missing boundaries in fact not being real boundaries, but rather incorrect calls which the simulations correctly fail to reproduce.

The 58 “extra” boundaries which are present in simulations but not found in the MicroC data were found to be flanked by nucleosomes depleted in most “active” histone modifications, but enriched in H3K36me3 (see Suppl. Fig. S3c). It was noted in Ref. [10] that H3K36me3 is highly depleted at MicroC boundaries – together with the results presented here this suggests that this mark is associated with a mechanism which promotes nucleosome-nucleosome interactions across a long linker (or NDR) which would otherwise act as a boundary. In yeast, di- and tri-methylation of H3K36 has been associated with transcription [26], and the Set2 enzyme responsible for generating these marks is thought to interact with RNA PolII in a way which is consistent with co-transcriptional H3K36 methylation [27]. While ~ 65% of MicroC boundaries within the simulated regions are at, or near to, gene promoters, this is only the case for ~ 9% of the “extra” boundaries. Together this suggests that transcription is another important determinant of boundaries: long linkers (which are normally associated with promoters) are natural boundaries, except when they occur within a gene body, in which case active transcriptional elongation appears to abrogate the boundary.

In order to compare our simulation results with the MicroC data at a level beyond domain boundaries, we look at how interactions depend on genomic distance. Figure 2c shows how the average interaction strength between two nucleosomes depends on their separation, *s*, within the simulated regions. From the experimental data we observe that on a log-log plot there appears to be two linear regimes indicative a power law relationship (mean number of reads ~ *s^−α^*): for short range (*s* ≈ 2–10 nucleosomes) α ≳ 2, while at longer range (*s* ≈ 10–100 nucleosomes) *α* ≈ 1. Similar power-law behaviour is observed in HiC data, and this has been used to infer a “fractal” organisation of chromosomes [28]. An equilibrium polymer coil is expected to display a power law scaling with exponent *α* = 3/2 for small *s* and *α* → 0 for large s, whereas HiC data often shows *α* ≈ 1 for all length scales [29]. It is interesting that the MicroC exponent differs from that of HiC at shorter genomic lengths: this may reflect the heterogeneity in fibre structure at this length scale. Our simulations are close to the MicroC data for short genomic separations, but show the equilibrium coil behaviour at longer range – this is to be expected since there is no component of the model which would lead to deviation from this (e.g., there is no long-range looping such as might be mediated by proteins *in vivo*). A discussion of the genome-wide scaling is given in the Supplementary Material (and see Suppl. Fig. S4-5).

### A more realistic nucleosome geometry *does not* improve domain predictions.

As noted above, it is surprising that our simple model can give such a good prediction of nucleosome interactions at the micro-domain level. We might expect that a more detailed representation of the nucleosome geometry, which is well known from crystallography [31, 32], may be an important aspect to include, and that it may increase the agreement with chromatin interaction data. We therefore now turn to a more sophisticated model (Fig. 3(a)) where: (i) we use a more realistic “disk-like” shape for the nucleosomes instead of a 10 nm sphere, and (ii) we simulate the way linker DNA enters/exits the nucleosome by including an angle constraint [19]. The more detailed description possesses some (but not all) of the features included in the highly detailed models described in Refs. [20, 21], which have been used in Monte Carlo simulations to study the folding of short arrays of regularly spaced nucleosomes into 30 nm fibres.

**Fig. 3:**
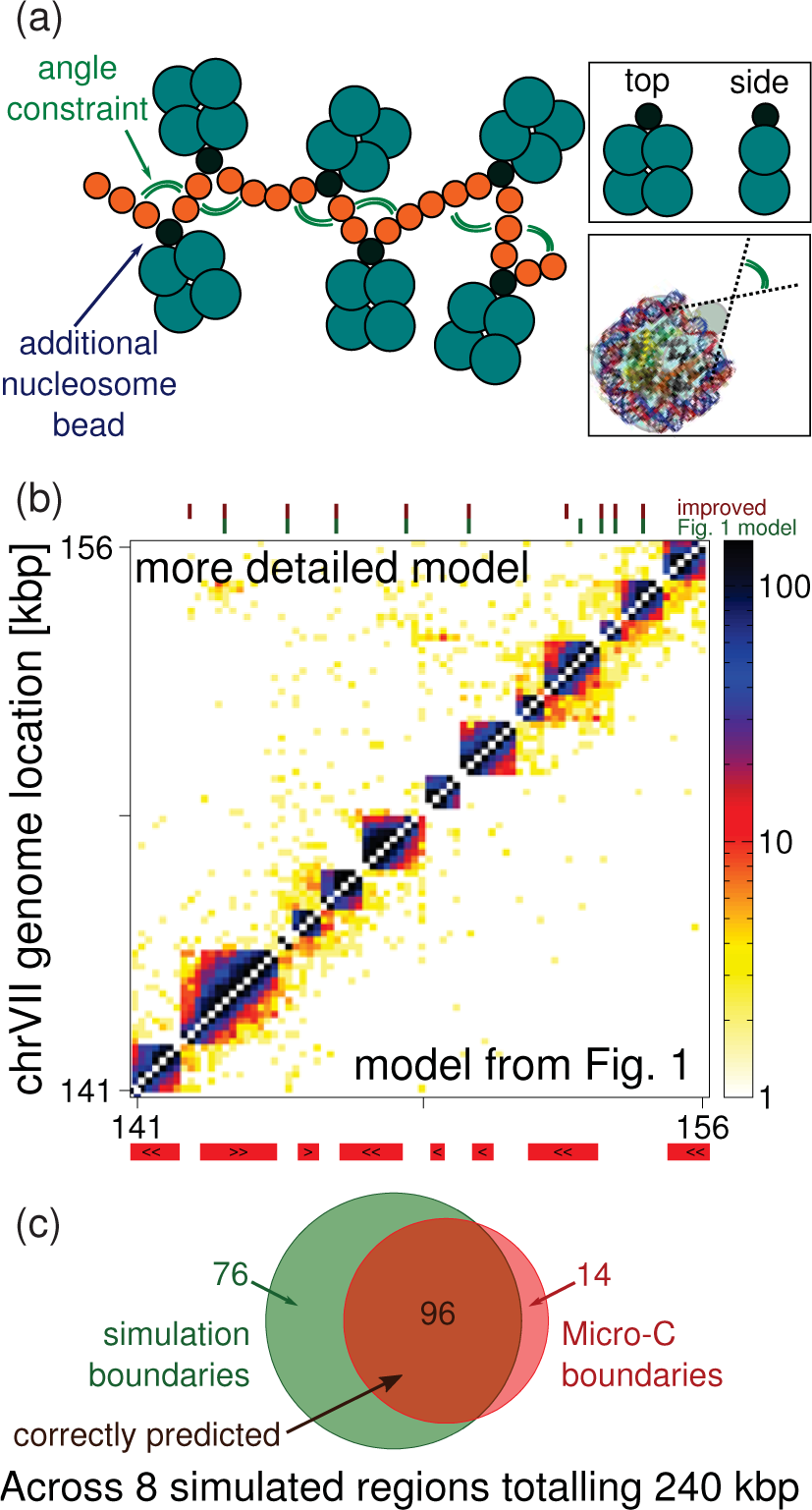
A more detailed model takes into account known aspects of nucleosome geometry. (**a)** Schematic showing the more detailed model. Nucleosomes are made of up from five beads, diffusing as a single object. Four larger beads arranged in a square approximate the disk shape of a nucleosome, with diameter roughly 10 nm and height 5 nm. The linker DNA is attached to the nucleosome through a smaller bead at one edge of the nucleosome, mimicking the way the entry/exit linkers leave the nucleosome on the same side at a preferred angle. Top inset: top and side views show a more disk-like nucleosome shape. Bottom inset: the nucleosome schematic is overlaid on an image of the nucleosome crystal structure (obtained from PDB: 1KX5, Ref. [30]) to show the preferred linker exit/entry angle. (**b)** Interaction map for region chrVII:140680-155644, where the upper half shows simulations using the model described in Fig. 1 and the lower half uses the more detailed model shown in panel a. (**c)** Venn diagram comparing boundaries predicted with the more detailed model, and those obtained from the MicroC data.

Intriguingly, despite the improved geometrical resolution of the nucleosomes, there is no appreciable improvement in agreement with the MicroC data. In Fig. 3b-c we show results for the version of the model where, as for the simpler model, there are no nucleosome-nucleosome interactions except for volume exclusion. Visually the interaction maps look very similar to those generated by the model of Fig. 1; likewise the agreement in boundary positioning shows little difference (over all simulated regions 87% of boundaries were predicted correctly, with 76 additional simulated boundaries); the correlation between the MicroC boundary signal and that from the new simulations is *r* = 0.42 (*p* < 10^−10^). Interestingly, however, there is a marked difference between the two models in terms of how the average interactions vary with separation (Suppl. Fig. S5) – at separation *s* > 20 nucleosomes the more detailed model deviates significantly from the MicroC data (see the Supplementary Material for further discussion).

Since the more complicated model does not show any significant improvement, for the rest of this work we return to the simpler model of Fig. 1. (Some further refinements which also fail to improve agreement with the data are described in the Supplementary Material.)

### Nucleosome spacing is irregular in yeast chromatin genome-wide

Visual inspection of the simulated chromatin conformations we generated (see Fig. 1b) shows that nucleosome spacing is highly irregular, and this leads to the formation of a heterogeneous fibre. Although nucleosome positioning data have been available for some time now, this fact is often overlooked in discussions of the formation of chromatin fibres (as typical textbook pictures usually show regular spacing). We now ask whether irregular nucleosome spacing is a generic feature of yeast chromatin *in vivo*, and we examine nucleosome spacing genome-wide.

Figure 4a shows the distribution of linker lengths across the 8 simulated regions; also shown is the genome-wide distribution (nucleosome positions generated using the NucPosSimulator software as before). First, we note the concordance between these distributions, indicating that the simulated regions are representative. Second, we note that the distribution is far from what would be expected for either regularly or randomly spaced nucleosomes. In the former case, one would expect a Gaussian distribution around a mean value; in the latter case, nucleosomes were positioned by a Poisson process, one would expect an exponential distribution. In fact, the distribution is multi-modal, with a large number of very short linkers (about 25% of linkers genome wide have length 1-3 bp) and a broad peak centred on ~16 bp. Interestingly there are also many linkers which are much longer (about 12% of linkers are between 50 and 200 bp), which presumably correspond to nucleosome depleted regions (NDRs), such as are found at gene promoters. (We assume most linkers greater than 200 bp are artefacts due to un-mappable regions of the genome.) Typically the nucleosome repeat length for yeast is quoted as 165 bp [34, 35], which corresponds to a linker length of 18 bp. From the distribution shown in the figure, the mean linker length is ~ 28.7 bp (and this decreases to ~ 18 bp if only those linker which are ≤ 100 bp are considered).

**Fig. 4:**
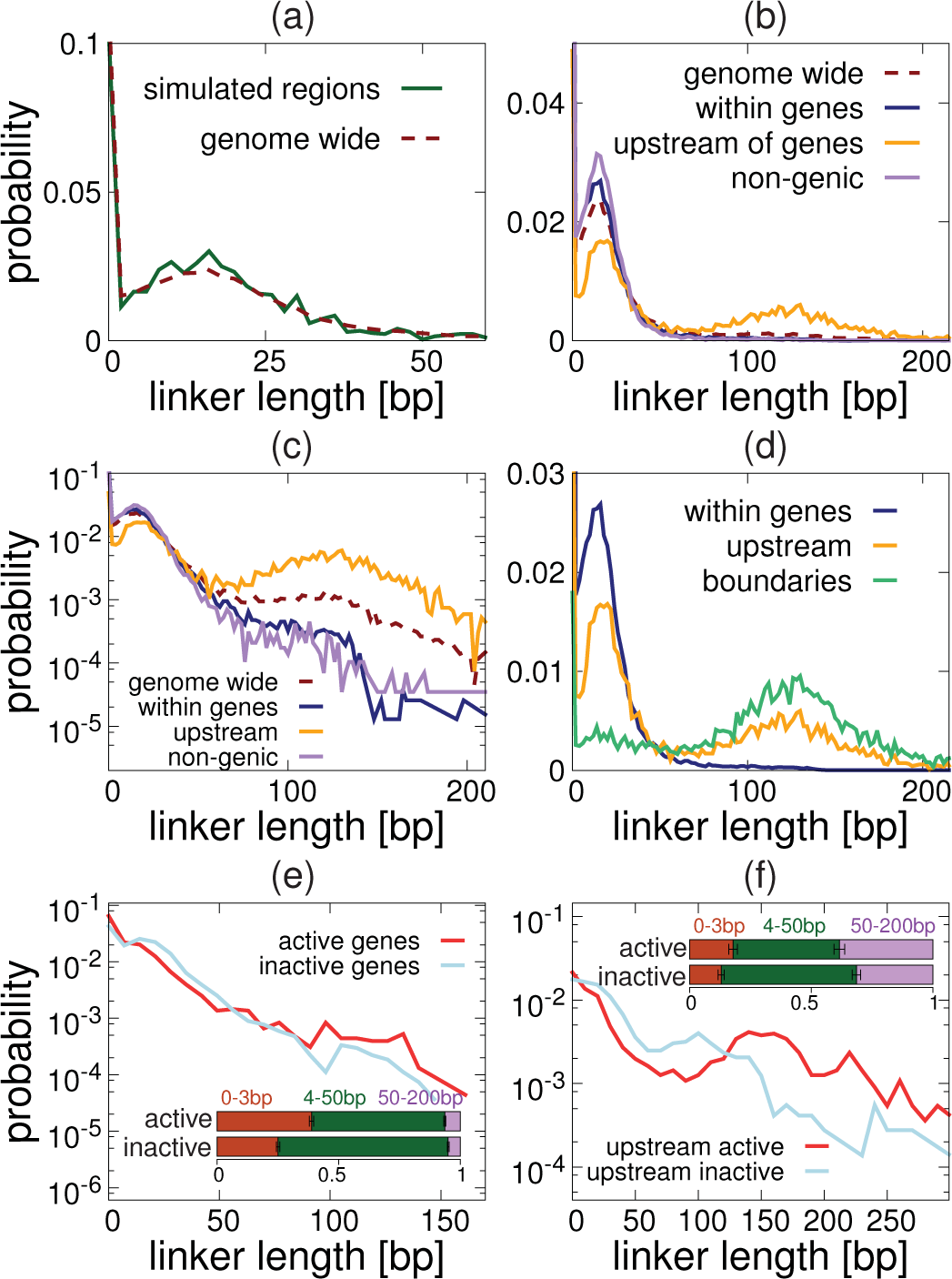
Linker lengths have a multi-modal distribution. Plots showing the linker lengths distribution based on nucleosome positions generated by NucPosSimulator (using MNase-seq data from Ref. [13]). (**a)** The genome wide distribution (red dashed line) is shown alongside that for the eight simulated regions (green solid line). (**b)** Separate distributions are shown for nucleosomes within genes (including all annotated genes of length ≥1 kbp), within the 500 bp upstream of gene transcription start sites (TSS; the same set of genes is used) and in non-genic regions of the genome. (**c)** The same plot as (b) is shown on log-linear axes. (**d)** The distribution of linker lengths found at domain boundaries (found genome-wide using the same method as described above), is shown alongside the ’within gene’ and ’upstream of gene’ distributions. (**e)** Distributions for nucleosomes within active and inactive genes are shown separately. (Activity is inferred from ChIP-on-chip data for PolII, obtained from Ref. [33], as detailed in the Supplementary Material). The inset shows the proportion of linkers with length less than 200 bp which fall into the three indicated length ranges, with error bars showing standard errors. Non-overlapping error bars indicate a statistically significant difference. (**f)** Similar plot to (e) but for linkers of nucleosomes in the 500 bp upstream of gene TSS.

In Figs. 4b-d we examine the linker length distribution more closely by separating out different types of genomic region. Specifically we look at linker lengths (i) within genes, (ii) within regions 500 bp upstream of genes (promoters)^2^, and (iii) in non-genic regions. In order to unambiguously identify linkers within gene bodies, we limit the analysis to genes of length ≥ 1 kbp in categories (i) and (ii), but consider all annotated genes when determining linkers in category (iii). We find that linkers within genes and in non-genic regions show a similar size distribution (though there are more short, < 3 bp linkers within gene bodies – ~ 30% compared to ~ 25%). As expected, in the promoter regions there are also many long (50-200 bp) linkers (~ 40%), and a lower proportion of short and medium length linkers. Figure 4d confirms that genome-wide, boundaries tend to be at long linkers (with the adjacent linkers tending to be short or medium in length).

In Figs. 4e-f we further separate active and inactive genes using PoIII binding data [33] as a proxy for transcriptional activity (we take genes with PoIII binding levels below and above the 10th and 90th percentiles respectively, see the Supplementary Material). This reveals that the bodies of active genes have more very short linkers, and fewer medium (4-50 bp) linkers than inactive genes (Fig. 4e). This is consistent with previous work [36] which found a correlation between gene activity and nucleosome density within coding regions (suggesting that, perhaps surprisingly, nucleosome crowding strongly facilitates transcription elongation), and proposed that transcriptional plasticity (the variation of gene expression as a result of environmental changes) may be facilitated by chromatin remodellers which alter nucleosome spacing. Other recent work [37] revealed a correlation between nucleosome crowding (i.e. closely spaced or even overlapping nucleosomes) and increased nucleosome turnover, which itself is associated with gene activity [38]. The promoter regions of the active genes also showed slightly more short linkers than their inactive counterparts, as well as more long linkers (Fig. 4f).

To check that the linker length distribution present in our simulation is not an artefact of the specific experimental technique used to generate the data (MNase-seq) or the simulated annealing process used to obtain nucleosome positions, in the the Supplementary Material we present a similar analysis of linker lengths obtained from site-directed DNA cleavage experiments, which offer higher resolution data than MNase-seq [39].

### Chromatin conformations with irregular (realistic) nucleosome spacing are heterogeneous and differ from regular fibres

We now use our simulations to examine some of the properties of 3-D structures formed by fibres with irregularly spaced nucleosomes, by comparing these to fibres of similar length with regularly spaced nucleosomes (Fig. 5a). First, we ask how nucleosome spacing affects the volume taken up by the chromatin fibre. Figure 5b shows how the radius of gyration (a measure of the size of the fibre) varies as a function of fibre length for the two cases. We calculate this by finding the *R_g_* of the first *L* beads of the fibre (treating DNA and nucleosome beads on the same footing), then beads 2-*L* + 1, then 3-*L*+2, and so on; we average over all such windows of length *L* and over snapshots taken at intervals during the simulation as before. We find that the irregularly spaced fibre is smaller than the regular case (*R_g_* reduces by about 10%); this could be interpreted as a decrease in the effective persistence length or stiffness of the fibre. Fitting a power law, we find a similar exponent in each case (*α* ≈ 0.64 for the irregularly spaced nucleosomes and *α* ≈ 0.67 for the regular case): these are likely finite *N* crossovers to the value expected for large *N* for a polymer in a good solvent (*α* ≈ 0.588).

**Fig. 5:**
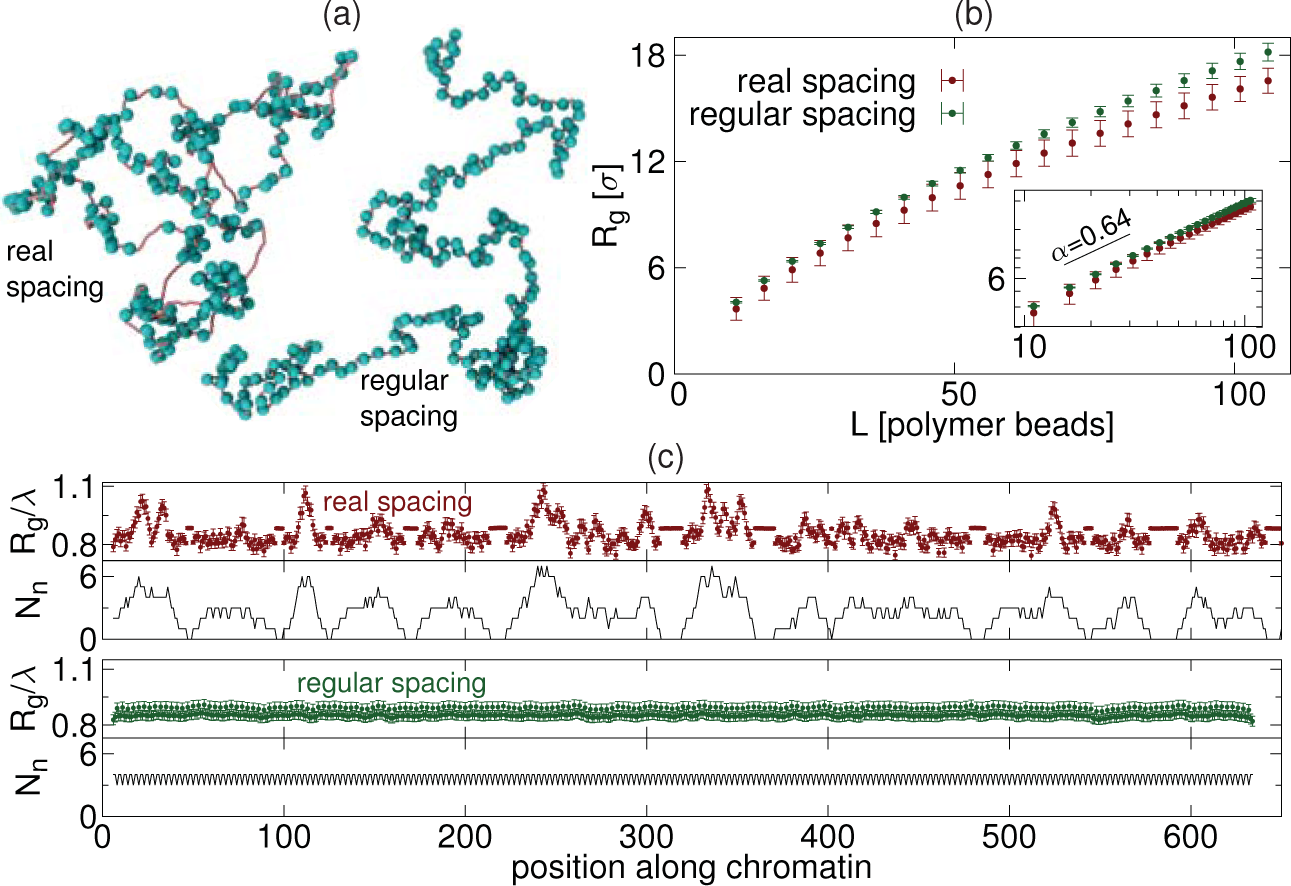
Irregular nucleosome spacing affects the local size and structure of the fibre. (**a)** Snapshots from simulations of: (left) yeast genomic region chrIV:1,254,937-1,287,938 using nucleosome spacing obtained from MNaseseq data (shown in Fig. 2c), and (right) a fibre of similar length with regularly spaced nucleosomes (linker length 22 bp). (**b)** Plot showing the radius of gyration, *R_g_*, as a function of the length of the polymer (measured in numbers of beads – see text). Lengths are given in units of *σ* = 2.5 nm. Error bars show the standard deviation. Irregular spacing tends to reduce the size of the polymer. Inset shows the same data on a log-log plot: a straight line indicates a power-law relationship (*R_g_* ~ *L^α^*). The black line shows the exponent obtained from a fit to the real nucleosome spacing case. (**c)** Top plots (coloured points) show the average *R_g_*/*λ* of an *L* = 11 bead region, as a function of position along the fibre, for the irregular (chrIV:1,254,937-1,287,938 region) and regular spaced cases. Here λ is the square root of the contour length within the window in simulation length units (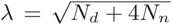). Error bars show the standard error in the mean. Bottom plots (black lines) show the number of nucleosomes *N_n_*. within the window.

Next, we examine how irregular nucleosome spacing affects the local fibre compaction, again using the radius of gyration as a measure. This time we use a fixed region length of *L* = 11 beads, and slide this window along the fibre, calculating at each position *R_g_* averaged over different snapshots and repeat simulations. Since the window consists of a mixture of DNA and nucleosome beads, we scale this by a factor 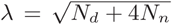, where *N_d_* and *N_n_* are the numbers of DNA and nucleosome beads within the window respectively (i.e. the square root of the contour length, since nucleosome beads have a size four times that of DNA beads). For the irregularly spaced fibre (Fig. 5c top) *R_g_*/*λ* varies widely with position along the fibre. The origin of the variation is revealed by a plot of the number of nucleosomes within each window (Fig. 5c, black lines). For a window with no nucleosomes we find *R_g_*/*λ* ≈ 0.88 ^3^. Adding a small number of nucleosomes to the region effectively introduces turning points into the polymer, and so reduces *R_g_*/*λ*. However, if many nucleosomes are added to the region the steric interaction between these leads to an effective stiffening of the chain and an increased *R_g_*/*λ*. For the regularly spaced nucleosomal fibre, the *R_g_* profile is, as expected, virtually flat.

These simulations show that different spacing of nucleosomes leads to relatively small, yet significant, differences in the global and local 3-D organisation of chromatin.

## Discussion

In this work we have presented a simple simulation model for chromatin, using it to study interactions within the chromosomes of the budding yeast *Saccharomyces cerevisiae*. Surprisingly this seemingly simple model, where nucleosomes are represented by 10 nm spheres connected by linker DNA (Fig. 1), had sufficient detail to correctly predict the nucleosome interaction patterns observed in recent MicroC data [10] (which revealed ‘micro-domains’ of typical length ~1-2 kbp). Specifically the simulations reproduced the drop off in the mean interaction frequency as a function of separation (at least for separations up to 10 nucleosomes), and were able to correctly determine the locations of 84% of micro-domain boundaries in 8 simulated genomic regions across 6 chromosomes. Additional microscopic details such as a more realistic nucleosome geometry, and constraints on the exit/entry angles for linker DNA were not required (i.e. a more detailed model including these features did not show any appreciable improvement in the agreement with the data).

Since the only data used as an input for the simulations was the genomic positions of nucleosomes, this implies that the pattern of micro-domain boundaries is largely encoded in these positions. While previous work [10] found domain boundaries to be enriched for binding of some proteins, and the flanking nucleosomes were enriched for transcriptional activating histone modifications, our results suggest that these are not directly responsible for boundary formation. Rather, protein binding (e.g., of chromatin remodellers) more likely gives rise to the formation or maintenance of nucleosome depleted regions, and this in turn forms a boundary. Our model also incorrectly predicted that boundaries would be present at some long linkers within gene bodies – this failure is however itself informative, since it suggests that transcription through NDRs can counter their boundary-forming potential.

In light of the important role of nucleosome spacing in chromatin interactions, we next examined the linker length distribution in more detail (using both positions generated from MNase data, and other experimental methods – see the Supplementary Material). The surprising number of very closely spaced or even overlapping nucleosomes, and the high abundance of these within (particularly the most active) gene bodies suggests that this has a role in transcription elongation [36].

Finally, we used our simulations to study how the irregularity of linker lengths affected the three-dimensional polymer properties of chromatin fibres. We found that chromatin fibres made from irregularly spaced nucleosome arrays leads to an overall reduction in the size of the polymer compared to a regularly spaced fibre of the same length. By examining the local compaction of a region of the chromatin fibre as a function of the position along it, we found that irregular spacing leads to wide variation of 3-D size compared to the regularly spaced nucleosomes case. In our model, this variation closely followed the number of nucleosomes within the region. Unexpectedly, we found a non-linear relationship between the number of nucleosomes within a region and its 3-D size – for a small number of nucleosomes the size reduces compared to a region with linker DNA only, whereas if many nucleosomes are present the 3-D size is larger. Since the entry/exit angle of the linker DNA (which is not constrained in our simple model) is also likely to have an effect, it would be of interest to study this with a more detailed simulation scheme in future.

In summary, our simulations have revealed a close link between nucleosome positioning and chromatin interactions in 3-D in yeast. Although genome-wide data on nucleosome positions have been available for several years, the striking irregularity in nucleosome spacing is often overlooked. It will be of interest to study how this affects the 3-D structure of chromosomes [40], and how chromatin might fold into higher order structures in more detail – for example future models could investigate the effect of the torsional rigidity of DNA, which controls the relationship between linker length and the relative orientation of adjacent nucleosomes. One must also bear in mind that there are still many challenges in obtaining nucleosome positions, and the maps generated to date rely on information form a population of cells – it is still unclear what the nucleosome landscape is like within a single cell [41].

In higher eukaryotes the family of H1 linker histone proteins, which have been shown to induce folding of regularly spaced nucleosomal arrays into 30 nm fibres *in vitro*, are highly abundant and found across the genome, particularly in heterochromatin (to the contrary, the yeast homologue *HHO1p* has been found not to be present through most of the genome, but rather only at restricted locations [42]). Our simulation snapshots showing irregularrly spaced nucleosomes are strikingly reminiscent of recent imaging experiments in human cells [43] which revealed spatially heterogeneous groups of nucleosomes known as “clutches”. It would be of interest to study irregular nucleosome spacing in that context – how it varies in different genomic regions, and what are the implications for higher-order fibre folding – particularly since H1 is thought to control nucleosome repeat length, and is found to be depleted near active genes and promoters. Similarly, it would be interesting to understand if linker length plays a role in domain boundary formation at larger length scales in higher organisms – although chromatin looping and interactions between regions with similar histone modifications have been implicated there [44], long linkers might lead to kinks or distortions in the chromatin fibre which promote certain loops, or they might provide a natural barrier to the (1-D and 3-D) spread of histone modifications [45]. This may provide a mechanical link from DNA sequence, through nucleosome positioning, to higher order chromosome organisation.

## Materials and Methods

In this work we perform Langevin dynamics simulation of a chromatin fibre modelled as a bead-and-spring polymer using the LAMMPS software [23]. In brief, a fibre which resolves individual nucleosomes is represented by a chain of two species of spherical beads. Small (2.5 nm diameter) beads represent linker DNA, while larger (10 nm diameter) beads represent nucleosomes. We use a common model for DNA [16–18] which correctly captures its bending stiffness, and includes steric interactions. LAMMPS integrates the Langevin equation for each bead in the simulation using a velocity-Verlet algorithm, where an implicit solvent provides a thermostat which results in a constant NVT ensemble. Full details of the model and simulation scheme are given in the Supplementary Material. For the simulations presented in Fig. 3 the nucleosomes are instead represented as a rigid body composed of five smaller beads, arranged to approximate a 10 nm × 5 nm disk, where linker DNA forms an entry/exit angle of 72°; again full details are given in the Supplementary Material.

We compare our simulations to MicroC data obtained from Ref. [10] (GEO:GSE68016); this is aligned to the *Saccharomyces cerevisiae* genome following the methods discussed in that reference. We further map the MicroC data onto the set of “most likely” nucleosome positions obtained from MNase-seq data (from Ref. [13], GEO:GSM53721) using the NucPosSimulator software [14], which uses a Metropolis Monte Carlo algorithm to position nucleosomes according to a potential landscape inferred from the MNase data. Full details are given in the Supplementary Material.

## Acknowledgements

We acknowledge the European Research Council for funding (Consolidator Grant THREEDCELLPHYSICS, Ref. 648050).

## Supplementary Material

Including six supplementary figures and supplementary text.

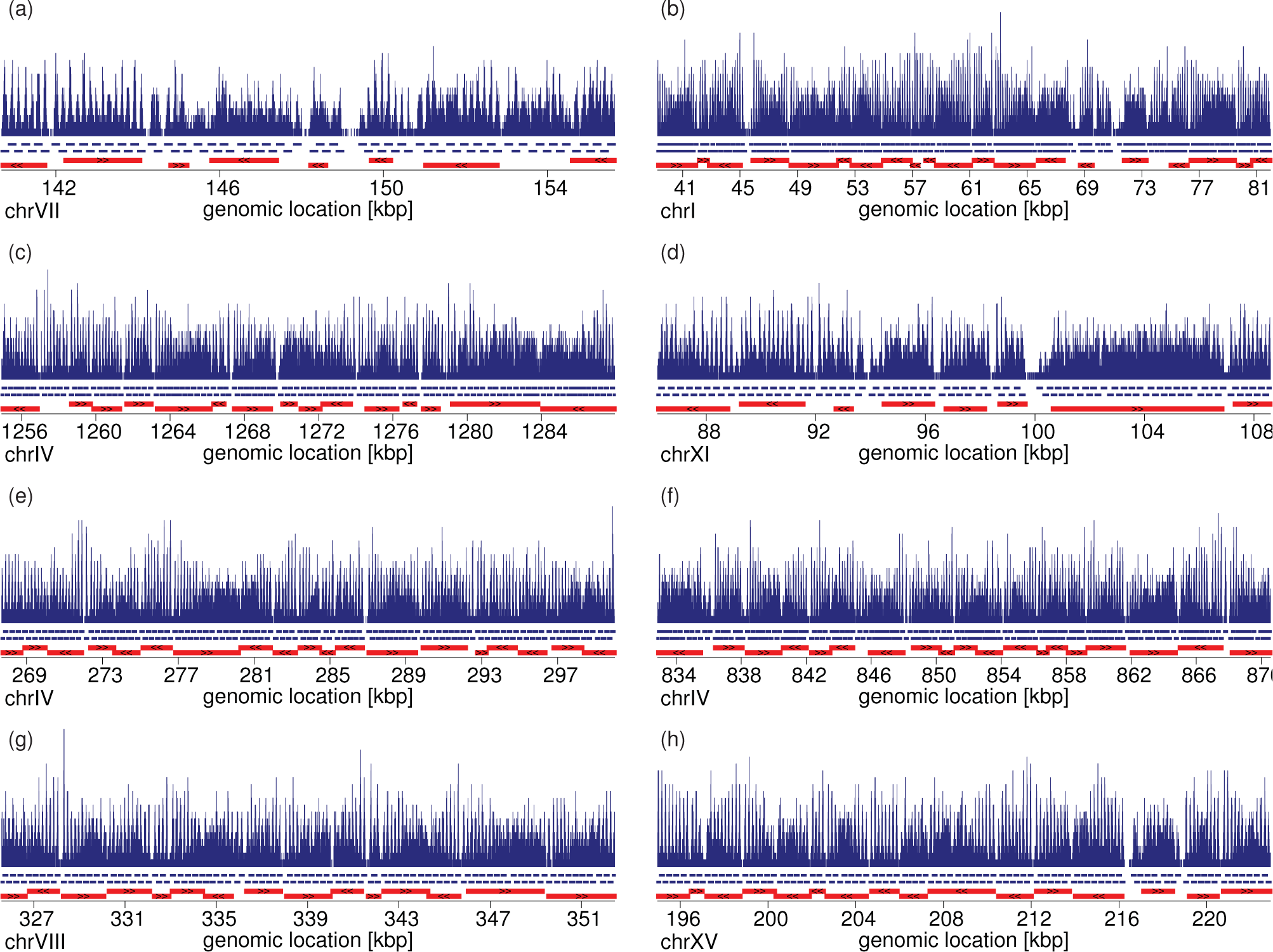
Suppl. Fig. S1: Plots showing the data used as an input for simulations for each simulated region. Top: pile-up of reads from yeast MNase data (from Ref. [1]) for the indicated genomic region. Blue lines under the pile-ups show the positions of nucleosomes as found using the NucPosSimulator software [2]. Red bars show genes (SacCer3 genome build).

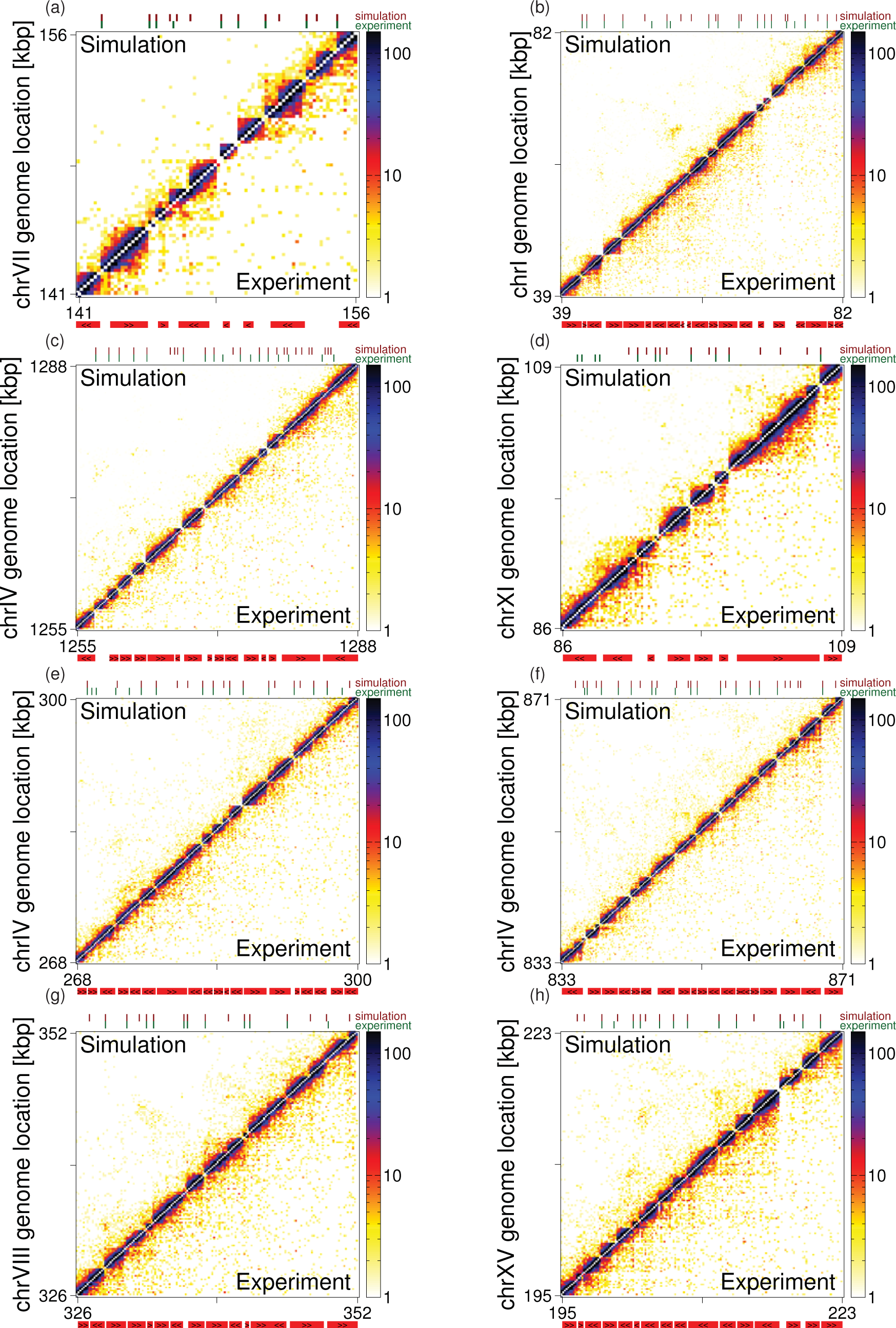
Suppl. Fig. S2: Maps showing interactions between nucleosomes in each simulated region. The colour of the square at coordinates *i, j* indicates the number of MicroC reads corresponding to interactions between nucleosomes *i* and *j*. The lower triangle shows MicroC data from Ref. [3], while the upper triangle shows simulation results. Numbers on the axes show genomic position, but these are approximate, since nucleosome spacing is not regular. The locations of genes (mapped to the nearest nucleosomes) are shown below the plot (red bars; gene orientation is indicated with black arrowheads). Domain boundaries were called from each map (see the Supplementary Material for details), and these are indicated with ticks above the plot (upper row shows simulation boundaries in red, lower row shows experimental boundaries in green).

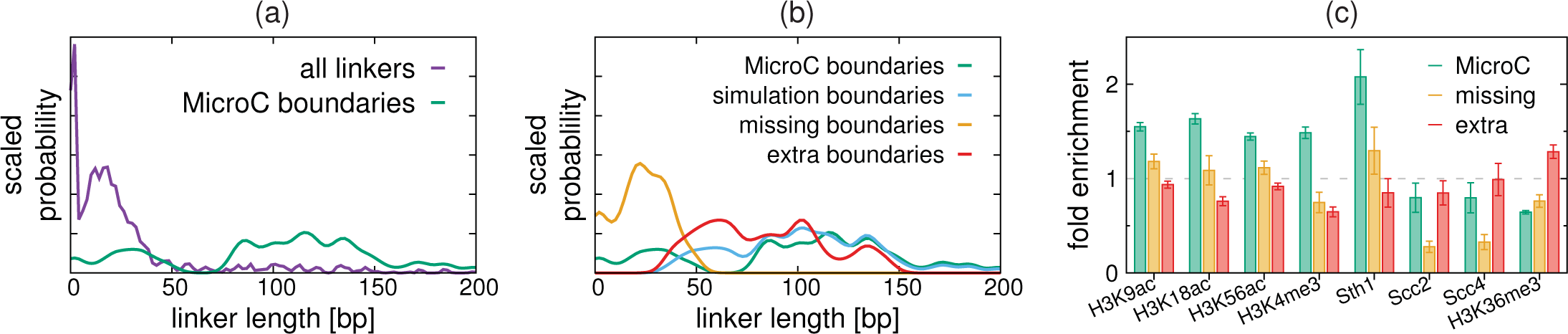
Suppl. Fig. S3: Plots showing the properties of boundaries within simulated regions. (a) The distribution of linker lengths across the simulated regions (purple) is plotted along side the distribution for linkers at boundaries called from MicroC data (green). A kernel density estimation method with bandwidth of 5 bp is is used, where curves are normalised to enclose unit area. (b) Similar distributions are shown for the boundaries found in simulations (blue), the “missing” boundaries (yellow; found in MicroC data but not predicted by simulations) and the “extra” boundaries (red; found in simulation but not present in MicroC data). Again curves are normalised to enclose unit area. (c) Mean fold enrichment of different histone modifications or protein binding levels are shown for the different classes of boundary. ChIP-seq data are obtained from Ref. [4] for histone modifications, and from Ref. [5] for protein binding. For histone marks fold-enrichment is against the ChIP input signal, whereas for proteins the fold-enrichment against the mean protein level across the simulated regions.

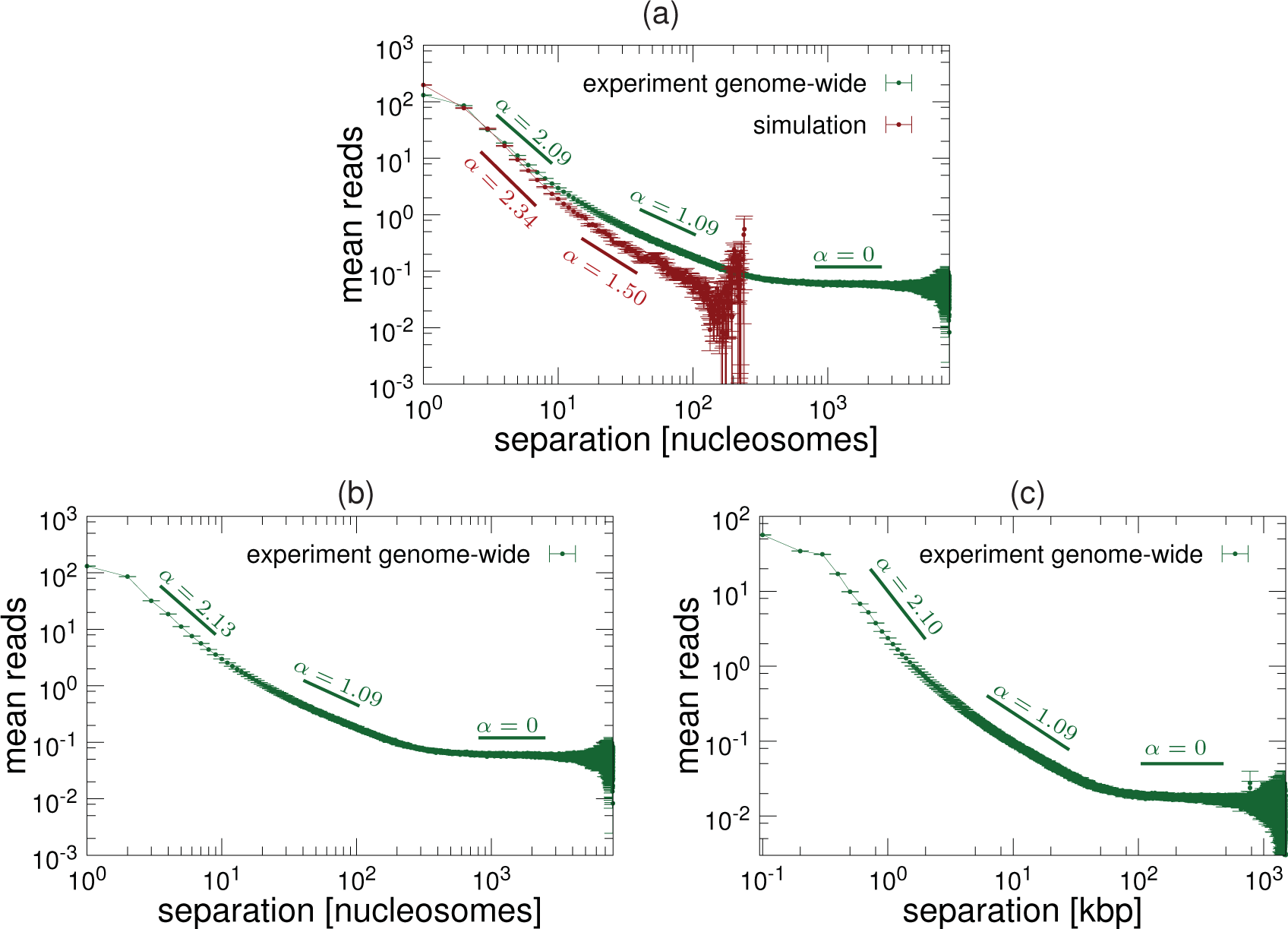
Suppl. Fig. S4: Plot showing how, on average, the number of interaction reads between nucleosomes scales with their genomic separation. A linear relationship on a log-log plot implies power law behaviour (read ~ *s*^−*α*^) and exponents *α* are approximated using linear fits to different ranges of the log-data. (a) Green points are from genome-wide MicroC data [3], red points from simulations. Separations are measured in nucleosomes (i.e., a separation of 1 means neighbouring nucleosomes). (b-c) The experimental genome-wide distribution is shown with separations measured in nucleosomes (b) and in DNA length (c). Note that only the exponent for short separations changes very slightly.

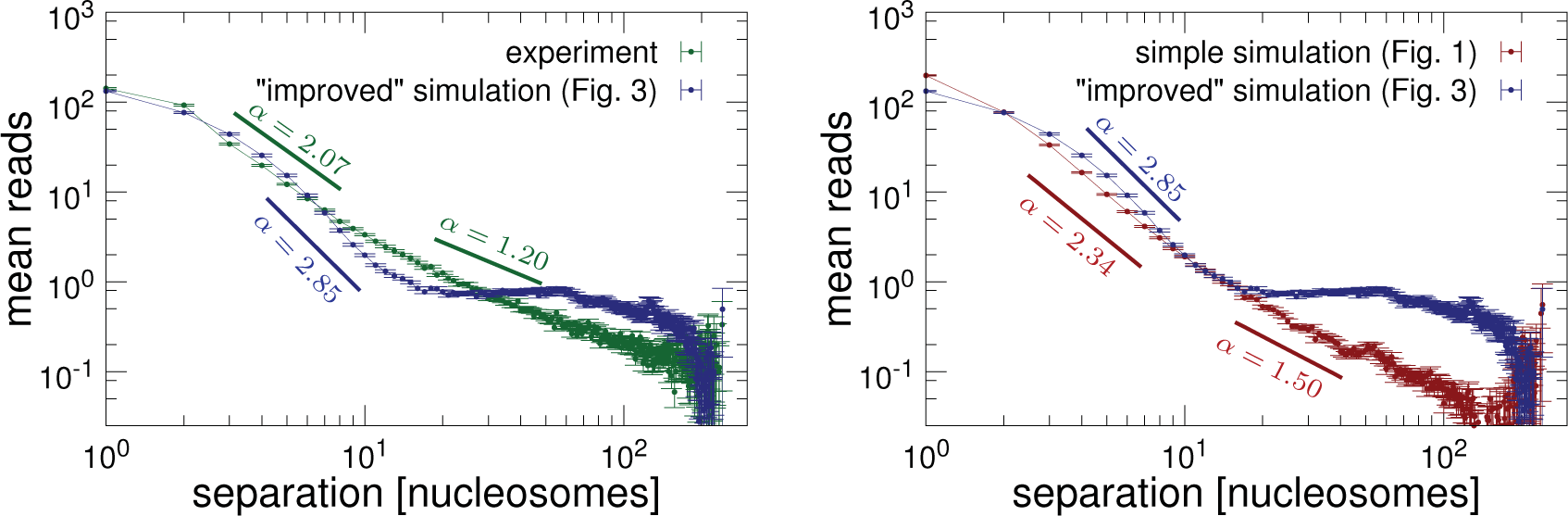
Suppl. Fig. S5: Plots showing how the number of interaction reads between nucleosomes scales with their genomic separation. (a) Comparing the simulation model shown in Fig. 3 with MicroC data. (b) Comparing the two simulation models.

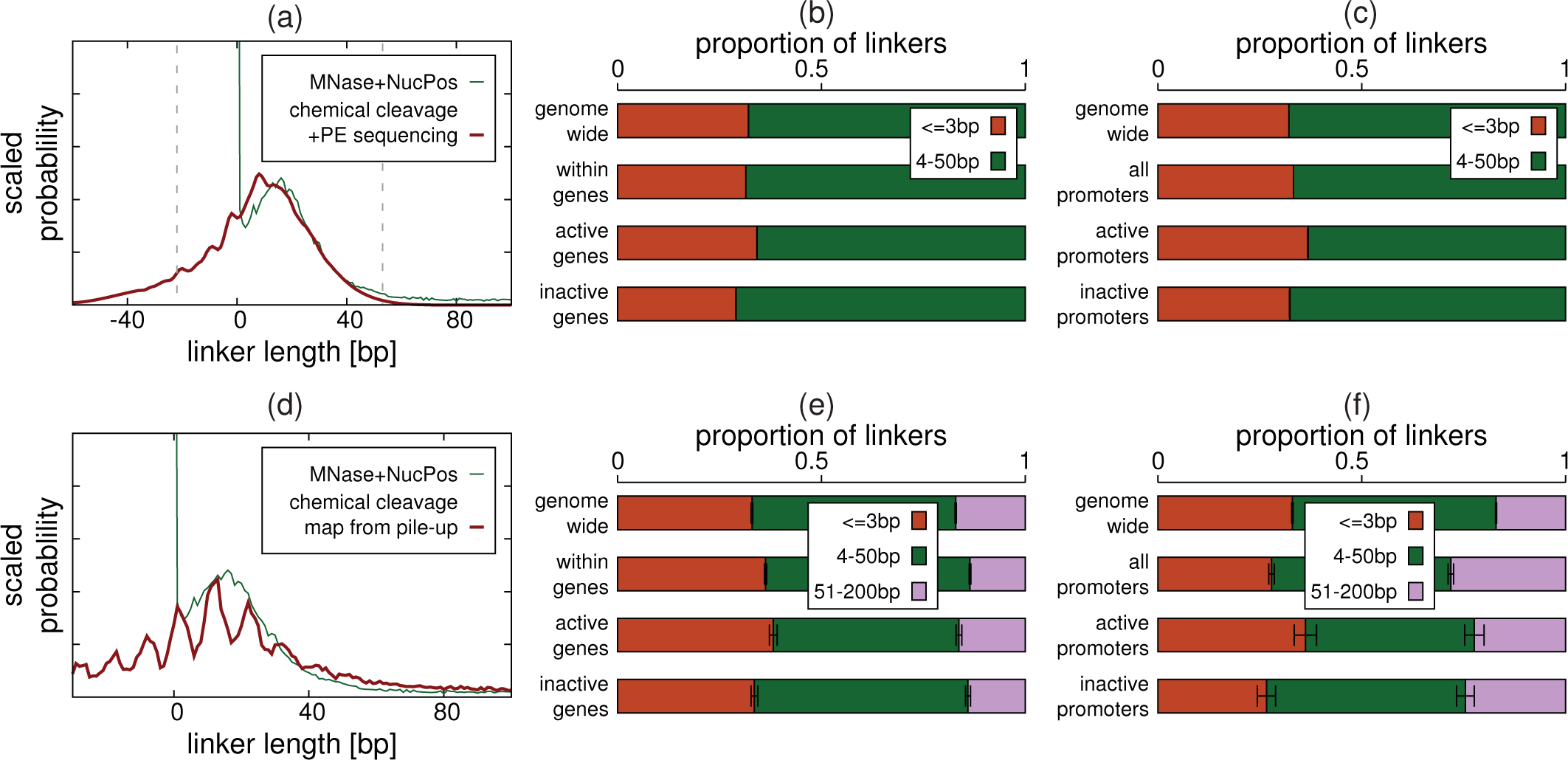
Suppl. Fig. S6: Plots showing linker lengths obtained from chemical cleavage data from Ref. [6]. (**a-c)** Plots of linker lengths obtained directly from chemical cleavage data using paired-end sequencing (see the Supplementary Material section *Genome-wide nucleosome spacing: further analysis*). The genome wide distribution of linker lengths (thick red line) is shown in (a), alongside the distribution obtained from MNase data using the NucPosSimulator software (thin green line; as shown in Fig. 4). Grey dashed lines enclose the range of linker lengths which are enriched by the experimental method. Linkers can be grouped into two size ranges, < 3 bp and 4-50 bp (with negative linkers referring to overlapping nucleosomes; this method biases against longer linkers), and by genomic location (within genes, or within the 500 bp upstream of genes) as discussed in the main text with reference to Fig. 4. Proportions of linkers falling within each group are shown in (b-c). Error bars are not shown as these are narrower than the lines. (**d-f)** Similar plots are shown for linker lengths based on a nucleosome positioning map obtained from pile-ups of cleavage sites. Here we use the map of “unique nucleosomes” (which forbids nucleosomes overlapping by more than 40 bp) which was provided as Supplementary Material in Ref. [6]. The strong ~ 10.5 bp periodicity identified in that reference is clearly visible. In (e-f) three linker size ranges are shown: < 3 bp, 4-50 bp, and 51-200 bp. The error in the proportions for the < 3 bp and 51-200 bp cases are shown as error bars; non-overlapping error bars imply that the difference is statistically significant. The same trends as in the MNase based map are observed.

### 1 Chromatin Model

Following previous work [7–9] we model linker DNA as a beadand-spring polymer, where beads of diameter 2.5 nm represent 7.35 bp of DNA. The *i*th bead in the chain, having position **r*_i_*** is connected to the *i* + 1th bead with a finitely extensible non-linear elastic (FENE) spring: the associated potential is given by

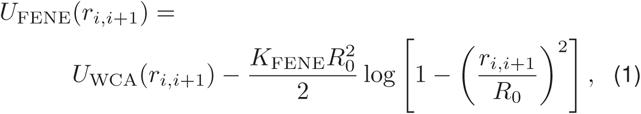

where *r_i,i_*_+1_ = |**r***_i_* − **r***_i_*_+1_| is the separation of the beads, and the first term is the Weeks-Chandler-Andersen (WCA) potential

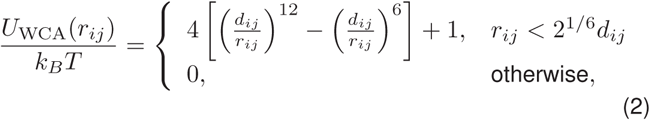

which represents a steric interaction preventing adjacent beads from overlapping. In Eq. (2) *d_ij_* is the mean of the diameters of beads *i* and *j*. The diameter of the DNA beads is a natural length scale with which to parametrize the system; we denote this by *σ*, and use this to measure all other length scales. The second term in Eq. (1) gives the maximum extension of the bond, *R*_0_; throughout this work we use *R*_0_ = 1.6 *σ*, and set the bond energy *K*_FENE_ = 30 *k_B_T* for linker DNA beads.

The bending rigidity of the polymer is introduced via a Kratky-Porod potential for every three adjacent DNA beads

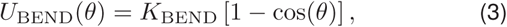

where *θ* is the angle between the three beads as given by

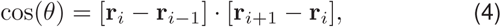

and *K*_BEND_ is the bending energy. The persistence length in units of *σ* is given by *l_p_* = *K*_BEND_/*k_B_T*.

Finally, steric interactions between non-adjacent DNA beads are also given by the WCA potential [Eq. (2)].

#### 1.1 Nucleosome Model of Fig. 1

In our first simple model, as depicted in Fig. 1a, nucleosomes are represented by 10 nm (4 *σ*) diameter beads, connected to linker DNA beads using FENE bonds according to Eq. (1) but with the appropriate choice for *R*_0_ (i.e. *R*_0_ = 3.6 *σ* for the bond between a DNA bead and a nucleosome bead, or *R*_0_ = 5.6 *σ* for the bond between two nucleosome beads); as before steric interactions between nucleosome beads, and between nucleosome and DNA beads are given by the WCA potential, with *d_ij_* being the mean of the diameters of the two beads.

#### 1.2 More Detailed Nucleosome Model of Fig. 3

In the more detailed model, as depicted in Fig. 3a, nucleosomes are represented by a rigid body composed of five component beads including four 5 nm (2 *σ*) beads and a 2.5 nm (*σ*) connector bead (see Fig. 3a). The four core beads are arranged with their centres on the corners of a square of size 4.2 nm (1.68 *σ*); the connector bead is positioned 5.75 nm (2.3 *σ*) from the centre of the square. Linker DNA beads are connected to nucleosome connector beads using harmonic springs with the associated potential

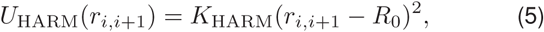

where *r_i,i_*_+1_ = |**r***_i_* − **r***_i_*_+1_| is the separation of the beads, and *R*_0_ = 1.1 *σ* is the equilibrium separation.

To constrain the entry-exit angle for linkers emerging from a nucleosome, a bending interaction between three connected DNAconnector-DNA beads is given by

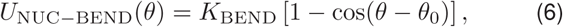

where *θ* is the angle between the three beads, and *θ*_0_ is the desired equilibrium angle, set at *θ*_0_ =72°, so as to match the entry-exit angle measured from the canonical nucleosome crystal structure [10]. We set the interaction energy to be the same as that used for the linker DNA beads.

### 2 Simulation Method

In our coarse grained molecular dynamics simulations, the position of the *i*th bead **r***_i_* changes in time according to the Langevin equation
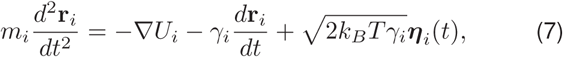

where *m_i_* is the mass of bead *i*, *γ_i_* is the friction it feels due to an implicit aqueous solvent, while ***η****_i_* is a vector representing random uncorrelated noise which obeys the following relations
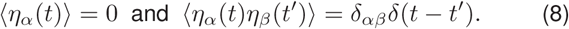

The noise variance is scaled by the thermal energy, given by the Boltzmann factor *k_B_* multiplied by the temperature of the system *T*, taken to be 310 K for a cell. The potential *U_i_* is a sum of interactions between bead *i* and all other beads, as described above. For simplicity we assume that all beads in the system have the same mass and friction *m_i_* ≡ *m*, and *γ_i_* ≡ *γ*. Eq. (7) is solved using the LAMMPS software [11] which uses a standard velocity-Verlet algorithm; we use a time step of Δ*t* = 0.005 *τ*.

### 3 Mapping simulation units to physical units

As detailed above our simulations use length units of *σ*=2.5 nm, and energy units of *k_B_T*; masses are given in units of the mass of a DNA bead, approximately 8 × 10^−24^ kg. A choice of *K*_BEND_ = 20 *k_B_T* therefore gives a realistic DNA persistence length of *lp* = 20 *σ* = 50 nm.

To map between simulation and real time units, we first note that the above defined length, energy and mass units lead to a natural time unit 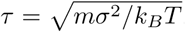. Another important time scale is the Brownian time *τ*_B_ = *σ*^2^/*D_i_*, which is the time scale over which a DNA bead diffuses across its own diameter *σ*. Here *D_i_* is the diffusion constant for bead *i*, given through the Einstein relation by *D_i_* = *k_B_T*/*γ_i_*. With our choice of *γ_i_* = 1 this means *τ*_B_ = *τ*. To map to real times we measure the mean squared displacement (MSD) for all beads; we then find the value of *τ*_B_

which gives the best fit to experimental results from Hajjoul et al. (2013), who measured the MSD for various chromatin loci in live yeast cells. This results in *τ*_B_ = 80 *µ*s, meaning that each 10^7^ time step simulation run represents approximately 4 s of real time.

### 4 Analysis of MNase data

Nucleosome positioning information is obtained from MNase-seq data published in Ref. [1] (available at GEO: GSM53721). Paired-end reads were aligned to the *S. Cerevisiae* reference genome (SacCer3 build) using bowtie2 [12]; duplicates and any reads with mapping quality less than 30 were removed. We then used the NucPosSimulator software [2] to obtain a map of the most likely nucleosome positions for each chromosome. In short, NucPosSimulator takes the centre point of each paired-end read and builds a frequency count profile, which is then used to generate an effective potential landscape for nucleosome binding. A Metropolis Monte-Carlo algorithm is then used to add, remove and move nucleosomes within this landscape. We use NucPosSimulator in “simulated annealing” mode, where the system starts at a high temperature (where nucleosomes will be highly dynamic) before being slowly cooled. During this process the nucleosomes settle into their most likely positions. Full details of the software are given in Ref. [2].

### 5 Analysis of Micro C data

MicroC data are obtained from Ref. [3] (available at GEO: GSE68016), and we follow a similar processing procedure as in that reference. After trimming adapters from the paired-end data we use bowtie2 [12] to align reads to the *S. Cerevisiae* reference genome (SacCer3 build); the data were treated as single-end reads, since ligation fragments will not form proper pairs when aligned. Duplicates were removed, and reads filtered to retain those where both pairs attained a mapping quality of at least 30. Following Ref. [2] we further filter reads according to the strand each read in the pair maps to, in order to avoid including reads resulting from runs of undigested nucleosomes. Interactions can then either be binned to obtain standard interaction maps, or can be mapped onto specific nucleosomes to obtain a nucleosome-nucleosome interaction map.

To generate nucleosome level interaction maps we use the nucleosome positions generated using NucPosSimulator and the MNase-seq data as detailed above. Treating each of the pair from each MicroC read separately, reads which overlap with a single nucleosome are unambiguous; reads which do not overlap with a nucleosome, but map to a position where their centre point is within 200 bp of the centre of one or more nucleosomes are assigned to their closest nucleosome. Reads which overlap with more than one nucleosome are assigned to that with which they have the largest overlap. Reads which do not map within 200 bp of a nucleosome, or which overlap with two nucleosomes by the same amount are discarded. Only read pairs where both members of the pair are assigned to nucleosomes are retained as informative interactions. Across 20 replicates, starting with 73,943,603 interactions, we were able to assign 73,803,602 of these unambiguously to pairs of nucleosomes; i.e. less than 1% of read pairs were discarded as it was ambiguous as to which nucleosomes they represented.

### 6 Generating Nucleosome interaction maps from simulations

From our simulations we obtain the positions of all beads in 2000 independent configurations for each region as detailed above. In a MicroC experiment nucleosomes are cross-linked, unprotected DNA is digested, and then protected DNA fragments are ligated: we expect that the probability that DNA from two nucleosomes are ligated together is a function of their 3-D separation. To mimic this process *in silico* we pick two nucleosomes at random from a simulated configuration, we then accept this as an interaction with a probability *P*(*r*) which is a function of the nucleosomes separation *r*, and reject it otherwise. For simplicity we choose a Gaussian shaped function 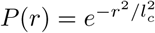, with interaction length scale *l_c_* = 15 nm (6 *σ*). For a simulation with *N* nucleosomes, we perform this operation *N*^2^ times. To obtain a number of simulated interactions which is similar to the number of MicroC reads for a given region we can repeat this entire process multiple times. Finally, to obtain a map which is comparable to the data, we scale all interaction counts by a factor *γ* which results in there being the same total number of reads in the simulated and experimental interaction maps.

### 7 Calling Boundaries from Interaction Maps

To find domain boundaries for both MicroC data and the simulated interaction maps we use a “sliding box” algorithm. Essentially a square box is placed off the diagonal of the interaction map (with its corner on *i*, *i*+1), and we sum the values for nucleosome interactions falling with the box. We then slide the box along the diagonal, nucleosome-by-nucleosome to obtain a boundary signal as a function of box position. Symbolically the signal at nucleosome *k* (used to determine if there is a boundary between nucleosomes *k* and *k* + 1) is given by

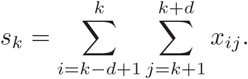

where we use *d* = 10. This is essentially the same as the algorithm used in Ref. [3], where for each nucleosome the number of upstream to downstream interactions (within some range) is counted. Minima in the signal indicate the positions of boundaries, and the value of the signal gives a measure of the boundary strength (the lower the value the stronger the boundary).

To identify boundaries we use the criteria that for position *k* to be a boundary we consider the value at the three upstream and three downstream neighbouring positions – five of the six values must be larger than that at *k*. This prevents small local minima from being falsely called as boundaries. We found that this procedure still gives some “false positive” boundary calls due to noise in the data (which is apparent when the maps are assessed visually); to remove these we take the distribution of the strengths of putative boundaries across the simulated regions, and discard any boundaries with a score above the 90th percentile of that distribution (higher value means weaker boundary).

We deem any boundary found in the simulated interaction map to be a “correct prediction” when it is within one nucleosome of the position of a MicroC boundary. As detailed in the main text, we denote a boundary which is found in the simulations but not the MicroC data an “extra boundary”, and any boundary found in the MicroC but not in the simulations is a “missing boundary”. In cases where two simulated boundaries are within one nucleosome of a MicroC boundary this is counted as one correct prediction, and one extra boundary; likewise if there is one simulation boundary within one nucleosome of two MicroC boundaries, this is counted as one correct prediction and one missing boundary.

### 8 Further model refinements also fail to show improved agreement with data

Given the surprising result that including a more realistic nucleosome geometry in the model does not give an improved agreement with the data, we attempted two further refinements. First we included a short ranged attractive interaction between nucleosomes to model direct nucleosome-nucleosome interactions which could be mediated by surface charges or histone tail interactions. We found no improvement in agreement with the data whether we included the attraction between all nucleosome, or only between a subset of nucleosomes. Since in Ref. [3] the authors noted that nucleosomes within inactive genes tended to interact more, we reasoned that the presence or lack of certain histone modifications could be used to identify nucleosome which might interact attractively – however this did not lead to an improvement in agreement with data. Second, we considered that certain histone modification might lead to partial unwrapping of DNA from the histone octomer. Although our model is too simple to include this in detail, we reasoned that it could be included in a simple way by removing the angle constraint on the entry/exit DNA for a subset of nucleosomes with acetylation modifications. Again we saw no improvement in agreement with the MicroC data.

### 9 Interactions *versus* genomic separation: genome-wide analysis

In the main text we discussed the mean number of interactions as a function of genomic separation (e.g., Fig. 2c), considering only the simulated regions. We noted that linear regions on a log-log plot implied a power law relationship, and that here there are two different linear regimes, in the 2-10 and 10-100 nucleosomes ranges. If we extend that analysis to the genome-wide case (Suppl. Fig. S4) we obtain similar exponents for those ranges, and observe the appearance of a new regime for *s* > 500 nucleosomes (roughly *s* >80 kbp). In this range the mean number of reads no longer varies with *s* (*α* = 0). It was already shown in Ref. [13] that the version of the MicroC protocol used to generate the data used here does not correctly capture these longer range interactions (the MicroC XL method gives data agreeing with the MicroC data at short separation, and HiC data [14] at longer range). We also note that, contrary to common HiC analyses, here we use coordinates of nucleosomes rather than base-pairs (Suppl. Fig. S4b-c shows that similar values of the exponents are obtained for both cases).

### 10 Genome-wide nucleosome spacing: further analysis

In the main text we present some analysis of nucleosome linker lengths based on MNase-seq data. We note here that similar results are obtained using data obtained from site-directed DNA cleavage experiments (from Ref. [6]), which offer higher resolution. Specifically the method uses an *S. Cerevisiae* mutant with a unique cysteine in histone H4, which allows chemical cleavage of DNA at precise locations near the nucleosome dyad.

Linker lengths can be obtained by two approaches. First, the fragments that the experiment yields are stretches of DNA between two nucleosome centres; these can be run on an agarose gel to reveal bands corresponding to DNA fragments from pairs, triplets etc. of nucleosomes. Purifying the lowest molecular weight band and performing paired end sequencing gives directly the separation of pairs of nucleosomes as they appear in single cells; however, this purification selects against longer linkers (fragments representing linker lengths between ~ –22 and 53 bp are enriched for; here a negative linker length means that the nucleosomes overlap). Suppl. Fig. S6a shows the linker length distribution obtained in this way, alongside that from the MNase data and NucPosSimulator discussed in the main text. Two features of note are that there are negative length values, which indicates overlapping nucleosomes, and that peaks at ~ 10 bp intervals are visible; we discuss these points in detail below. Though we cannot use the chemical cleavage data to analyse long linkers, we can still count the number short (< 3 bp) and medium (4-50 bp) length linkers in different genomic regions. Suppl. Fig. S6b-c shows that, consistent with the MNase based positioning (Figs. 4e-f), both within gene bodies and within the region upstream of their TSS, the proportion of linkers which are shorter than 3 bp is higher for active than for inactive genes. The fact that a similar distribution, and similar trends for different genomic regions, can be observed implies that the features discussed in the main text are not artefacts arising because nucleosome positions are based on a population level map.

The second approach to obtaining linker lengths from the chemical cleavage data, as detailed in Ref. [6], is to pile-up the strand-dependent cleavage points across the genome to form a nucleosome position map. Due to the specific pattern of cleavage positions on opposite strands, nucleosome centres can be identified from the strand-specific shape of the peaks (since the method is sensitive to the shape of the peak and not the height, the purification bias discussed above does not have an effect [6]). Here we used the “unique nucleosome map” provided as supplementary material in Ref. [6]; this does not allow nucleosomes to overlap by more than 40 bp (taking the positions with the highest score where larger overlaps occur). We note that, like MNase data, this map gives a “population” picture, and may not represent the situation within individual single cells. From this map we obtain the proportions of short (< 3 bp), medium (4-50 bp), and long (51-200 bp) linkers as in Fig. 4, and again find that short linkers appear more frequently within active than inactive genes and promoter regions (Suppl. Fig. S6e-f), and that long linkers (NDRs) appear more frequently in promoter regions than genome wide (though the difference between active and inactive promoters was not statistically significant, Suppl. Fig. S6f).

As detailed in Ref. [6], the higher-resolution obtained by the chemical cleavage method reveals that a significant proportion of nucleosome overlap with the territory of their neighbours, and that there is a strong ~ 10.5 bp periodicity in the linker length distribution. The former manifests itself in the data as nucleosome centre-centre DNA fragments shorter than 147 bp (negative linker lengths); this has been studied using an *in vitro* positioning system [15] – there the authors observed that partial unwrapping of DNA from the histone octomer allows nucleosomes to invade their neighbour’s territory and that extreme overlapping coincides with a loss of an H2A-H2B dimer from one nucleosome. In Ref. [16] a statistical mechanics model which constructed a nucleosome-DNA binding free energy based on the assumption of a strong DNA-histone interaction at points where the minor groove contacts the protein could reproduce the observed distribution of (positive and negative) linker lengths – as DNA unwraps there is an energy increase because hydrogen bonds are broken, but at the same time the entropy of the unwrapped section increases, leading to an oscillatory free energy. This model (which does not have a sequence dependent component) also explains the linker length periodicity through a DNA-nucleosome free energy profile which extends beyond the 147 bp footprint – this could arise through DNA-histone tail interactions, steric interactions between neighbouring nucleosomes (the bp separation of nucleosomes determines their relative orientation), or interactions with other proteins such as H1 linker histone (HHO1p in yeast). Intriguingly if we use the NucPosSimulator software with the chemical cleavage pile-up data as an input to obtain the most likely nucleosome positions, the linker periodicity is lost; this is despite the other reported trends being retained, and the positions of over 80% nucleosome centres being with 40 bp of those of the “unique map” obtained from peak finding. NucPosSimulator assumes a “hard” steric nucleosome-nucleosome interaction, and gives similar linker distributions even if the nucleosome footprint is reduced to allow overlaps; this suggests that a “soft” nucleosome-nucleosome interaction would be required to retain the linker periodicity (as implied by Ref. [16]). At any rate, a highly irregular pattern of linker lengths is prevalent across all of these nucleosome mapping methods.

## Footnotes

1 The data presented in Ref. [10] did not show the long range interactions (particularly centromere-centromere and telomere-telomere interactions) observed in earlier low-resolution studies [15]. Improvements to the MicroC method, including the use of different agents for fixation [11], led to data which shows both the high resolution, shorter range interaction patterns and the longer range interactions. Since the original MicroC [10] study provided larger data sets, and here we are anyway interested in short range interactions at the micro-domain level, we use data from that work.

2 Note that in some cases the region 500 bp upstream of a gene overlaps with the 3′ end of the adjacent gene on the same strand, or the promoter region of an adjacent divergent gene.

3 The persistence length for our model DNA is 20 beads, so following the worm like chain model in the rigid rod limit we would expect a value not far from the upper bound 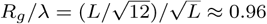, in agreement with our data.

